# 3D volumetric muscle reconstruction of the *Australopithecus afarensis* pelvis and limb, with estimations of limb leverage

**DOI:** 10.1101/2022.11.24.517817

**Authors:** Ashleigh L. A. Wiseman

**Affiliations:** McDonald Institute for Archaeological Research, University of Cambridge, UK

**Keywords:** *Australopithecus afarensis*, biomechanics, moment arm, muscle modelling, musculoskeletal

## Abstract

To understand how an extinct species may have moved, we first need to reconstruct the missing soft tissues of the skeleton which rarely preserve, with an understanding of segmental volume and the muscular composition within the body. The *Australopithecus afarensis* specimen AL 288-1 is one of the most complete hominin skeletons. Whilst it is generally accepted that this species walked with an erect limb, the frequency and efficiency of such movement is still debated. Here, 36 muscles of the pelvis and lower limb were reconstructed in the specimen AL 288-1 using 3D polygonal modelling which was guided by imaging scan data and muscle scarring. Reconstructed muscle masses and configurations guided biomechanical modelling of the lower limb in comparison to a modern human. Muscle moment arms were calculated and summed per muscle group. Simulated error margins were computed using Monte Carlo analyses. Results show that the moment arms of both species were comparable, hinting towards similar limb functionality. Moving forward, the polygonal muscle modelling approach has demonstrated promise for reconstructing the soft tissues of hominins and providing information on muscle configuration and space filling. This approach is recommended for future studies aiming to model musculature in extinct taxa.

## 1.0 Introduction

Soft tissues rarely preserve in the fossil record, rather we are mostly left with just the skeletal material. Yet, muscles animate the body. They allow an animal to move, walk and run. To understand how an extinct species may have moved, we first need to reconstruct the missing soft tissues of the skeleton with an understanding of volume and the composition within the body. Whilst these challenges in reconstructing hominin musculature have been addressed to an extent in the past (Berge, 1994; Karakostis et al., 2021; Kozma et al., 2018; McHenry, 1975; Nagano et al., 2005; Pontzer et al., 2009; Sellers et al., 2005; Wang et al., 2004), prior studies have typically relied upon estimated attachment sites consisting of a singular landmark and a simple action line representing leverage of the muscle (e.g., McHenry 1975; although see Karakostis et al., 2021). However, with the development of advanced computational methods, more sophisticated muscle modelling techniques have advanced musculature estimates of functionality (e.g., Nagano et al. 2005).

Although more recent methods are still founded in modelling lines of action which represent a muscle’s volume and path (Seth et al., 2018; Seth et al., 2011), these action lines are more sophisticated than earlier studies (e.g., Berge, 1994) by the utilisation of *via points* and *wrapping surfaces* (Carbone et al., 2015; Modenese & Kohout, 2020). These computationally advanced models produce well-validated estimates of limb functionality and leverage of a given extant species which have proven extremely informative (Charles et al., 2016; Hutchinson et al., 2015; van Beesel et al., 2021; Wiseman et al., 2021), but such modelling relies upon *known* muscle configurations within a limb, which is obviously problematic for musculoskeletal modelling of extinct species (e.g., Bishop et al. 2021 Demuth et al. 2022). Previous studies which have utilised sensitivity analyses during the muscle modelling process have demonstrated that changes in attachment site and path throughout a body can influence outputs (Charles et al., 2016; Hutchinson et al., 2005; Modenese & Kohout, 2020; O’Neill et al., 2013; Regnault & Pierce, 2018; Wiseman et al., 2021). Therefore, it is preferential for biomechanical models to be subject-specific and founded in known muscular compositions obtained via imaging methods (Carbone et al., 2015; Charles et al., 2020; Modenese & Kohout, 2020), albeit this is a momentous and challenging task for modelling the musculature of an extinct species in which this is *unknown*.

To be able to elucidate the mechanics which facilitate movement in fossil species, we first must have a comprehensive understanding of the un-preserved musculature, including surface attachment sites, volume and configuration (i.e., ‘this muscle’s belly overlays that muscle’s belly’). Such a challenging task was recently addressed by Demuth and colleagues (2022) in which 3D muscles were digitally reconstructed for an extinct archosaur and then their new method was validated with data from comparative extant archosaurs and then for mammals (specifically, primates) using polygonal modelling. In this approach, a 3D model of each muscle in a body segment is created, guided by comparative data from analogous extant species. The respective entire musculature configuration within an animal’s body segment is composed, whilst the entirety of the attachment site is digitised, not just a singular landmark. After which, a line is automatically ‘threaded’ through the midline of each newly created 3D muscle, representing each muscle’s line of action (LoA) which is appropriately configured and spaced within the limb. The method was successful with high fidelity, shows great promise and usability, and follows a simple workflow for implementation in future studies focused on hominins – a topic of great functional debate.

The *Australopithecus afarensis* specimen AL 288-1 (commonly known as ‘Lucy’), dated to 3.2 million years ago (Ma) from the Hadar region of Ethiopia (Johanson et al., 1982; Kimbel et al., 1994), is one of the most complete hominin skeletons and it has been well-studied since its discovery in the 1970s (Johanson et al., 1982; Nagano et al., 2005; Sellers et al., 2005; Tague & Lovejoy, 1986; Wang et al., 2004). AL 288-1 is predicted to have been 1.05 m tall (Jungers, 1982) with a body mass range of 13 to 42 Kg (Brassey et al., 2018; Grabowski et al., 2015; Ruff, 2010). This specimen is typically considered to be on the lower end of the body mass spectrum of the species (Jungers et al., 2016). Today, it is generally agreed amongst researchers that the postcranial skeleton displays morphological features indicative of bipedality (Gruss et al., 2017; Lovejoy, 1975, 2005, 2007; Ward, 2002), and has thus been the focus of previous biomechanical assessments investigating musculature capability (Nagano et al., 2005; Sellers et al., 2005; Wang et al., 2004).

The AL 288-1 specimen exhibits many anatomical features that differ to humans (Johanson et al., 1982), but typically the pelvis and lower limb (henceforth referred to as just ‘limb’ for brevity) anatomies have received more direct research attention as these elements are directly correlated with this species’ locomotory functionality. Such anatomical differences have polarised previous debates regarding the capability of *Au. afarensis* to walk bipedally with an erect limb (Rak, 1991; Senut, 1980; Stern, 2000; Stern & Susman, 1983). AL 288-1 has a wider pelvis and relatively shorter legs than humans (Johanson et al., 1982; Lovejoy, 1975; Tague & Lovejoy, 1986), which is thought to have effected muscular leverage (Berge, 1994; Pontzer et al., 2009; Sockol et al., 2007).

Whilst it is generally accepted today that AL 288-1 likely walked with an erect limb as humans do, questions still persist regarding the capability and frequency of such movement. Computer-based three-dimensional musculoskeletal models can be used to compute muscle function (Delp et al., 2007; Hutchinson et al., 2015; Wiseman et al., 2021), such as the calculation of moment arms (Charles et al., 2016; Seth et al., 2018; van Beesel et al., 2021). A muscle’s moment arm is defined as the perpendicular distance between the LoA and the joint axis, representing the effectiveness with which the force produced by a muscle generates moments (or torques) at the joint(s) crossed by the muscle. Moment arms, whilst informative, tell only a part of the story. Moment arm magnitudes do no correlate with the actual moment that a muscle can produce and, as such, magnitude alone should not be used to infer locomotor capabilities (Pandy, 1999). Patterns and peaks of moment arms are more informative and provide a baseline for inferring similarities in limb functionality between species.

Moment arms are intrinsically linked to bone shape, size and muscular composition (Pandy, 1999), and thus care must be taken in defining a muscle’s LoA in fossil specimens for which no musculature data exists. Changes/inaccuracies in the path can produce incorrect moment arm calculations (Brassey et al., 2017), necessitating new approaches beyond the use of identifying a simple straight-line LoA between two landmarks (e.g., Berge et al. 1994). Rather, configuration (i.e., each muscle’s volume and position within a body segment relative to other muscles) must be estimated to permit improved identification of attachment sites and LoAs.

By first reconstructing soft tissue composition in the body, more accurate estimates of each muscle’s LoA can be generated, producing realistic results of muscle function which are phylogenetically informed – that is, the researcher must first consider the most analogous extant species to inform upon the condition for an extinct specimen. In this study, soft tissues of the entire pelvis and limb of AL 288-1 were reconstructed using a digital polygonal muscle modelling approach which guided musculoskeletal modelling of this specimen to demonstrate the applicability of this approach. This study aims to provide a comprehensive production of muscle space filling and path designation in AL 288-1 and to make all reconstructions available for future use so that others interested in musculoskeletal modelling of hominins may use this model as a basis.

## 2.0 Materials and Methods

### 2.1 Human Data

A modern human (henceforth, just ‘human’) was selected as the closest probable analogy to the AL 288-1 specimen due to many anatomical similarities. MRI scan data of a human’s pelvis and lower limbs were sourced from an open-access repository (specimen ID: Subject03) (Charles et al., 2020). Subject03 was an adult female, weighing 72.6 kg and 176 cm in height. All extrinsic and many – but not all (see below) – intrinsic muscles were previously segmented and available for use. Segmental mass properties were readily available for this specimen. Unfortunately, the scan quality was too poor to conduct further segmentation to capture each muscle’s tendon (and thus attachment sites) and smaller intrinsic muscles, such as the hip external rotator compartment (i.e., *M. glemellus* group). For the former, each muscle’s insertion and origin were estimated, guided by muscle scarring and published dissection data for the right limb (Sedlmayr et al., 2022). Dissection data guided the latter to produce estimated muscle paths. A LoA was created for each muscle (Figure 1). All right muscle LoAs were mirrored to the left side, using the central location of the pelvis (location: 0,0,0) (Gatesy et al., 2022; Wiseman et al., 2022) – described below.

**Figure 1.**
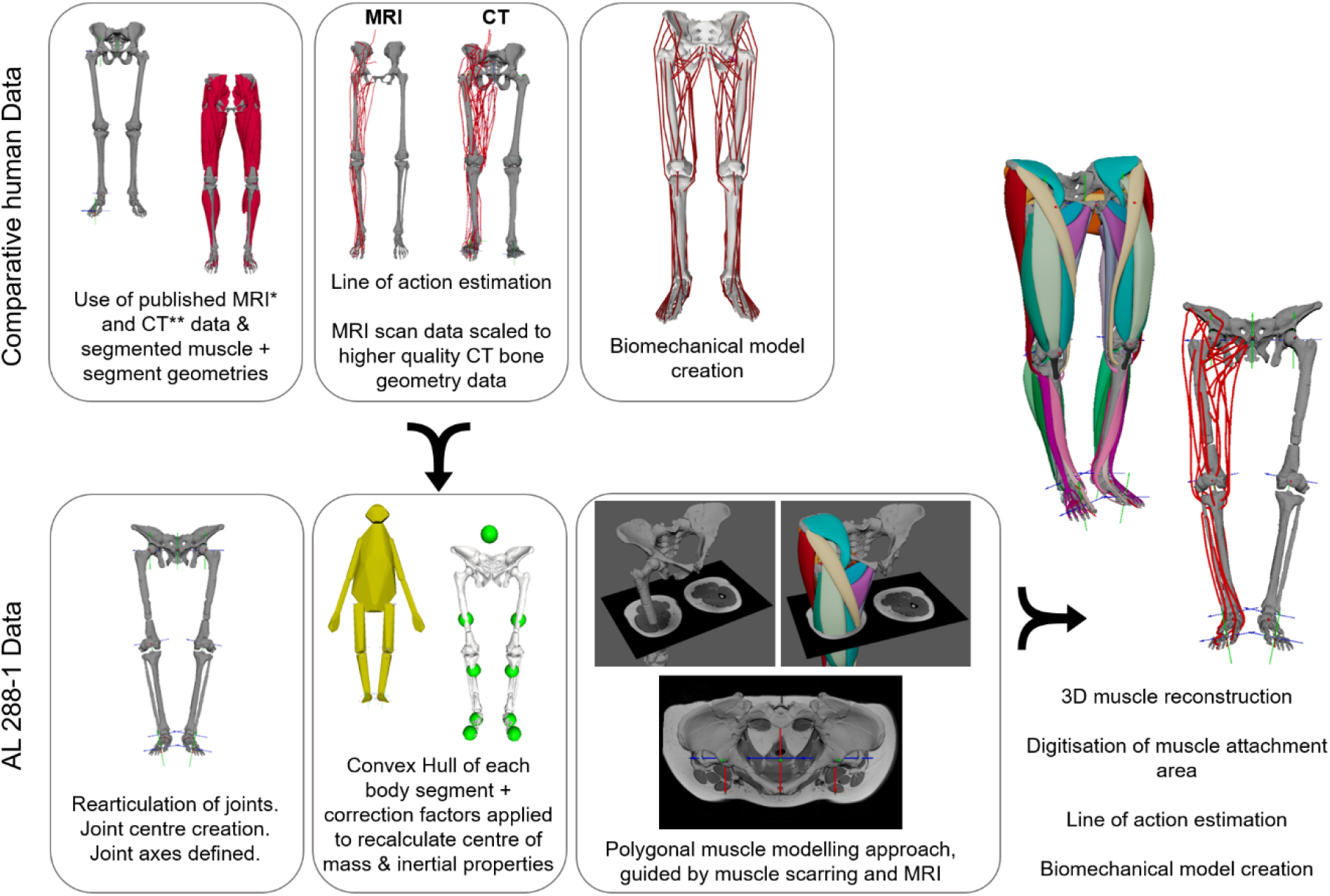
Workflow diagram outlining the process for the current study. Comparative data was collected from a male human, in which MRI data from (Charles et al., 2020) was used to guide muscle LoAs, segmental and inertial properties. The bone geometries were poor quality, necessitating these parameters to be scaled to high quality CT data from (Carbone et al., 2015; Modenese & Renault, 2021). For both the human and AL 288-1, joint centres and axes were created. A convex hull was created of each body segment of AL 288-1, of which the centre of mass and inertial properties were corrected following Coatham and colleagues (2021). A polygonal muscle approach was implemented (Demuth et al., 2022), producing muscle LoAs which were implemented into the biomechanical model of AL 288-1. Note: whilst (Charles et al., 2020) included the *M. psoas major* (visualised here), it was excluded from the current study due to uncertainty over thorax reconstructions.

The bone geometries of this specimen were low quality, and thus higher quality bone geometries of an adult male were sourced from other studies to better visualise bone epiphyses for the shape fitting procedure (see below), specimen ID: TLEM2-CT (Carbone et al., 2015; Modenese & Renault, 2021). This specimen weighed 84 Kg, height was not recorded.

Using the bone geometries from TLEM2-CT, joint centres and anatomical coordinates systems (ACS) were created, implemented via a shape fitting procedure (Bishop et al., 2021; Gatesy et al., 2022; Kambic et al., 2017a; Wiseman et al., 2021; Wiseman et al., 2022). Further details are provided in SI1. ACSs were established for the pelvis, hip, knee, ankle (the talocrural joint) and metarsophalangeal (MTP; other foot joints not modelled) joints (Gatesy et al., 2022; Kambic et al., 2014, 2017b). The hip was permitted to move along three degrees of freedom (DOF): flexion-extension, adduction-abduction and internal-external rotation; no translational DOFs were permitted (Wiseman et al. 2022). The knee, ankle and MTP joints were permitted to move along just one axis: flexion-extension. The X axis was abduction/adduction, Y was long-axis rotation, and Z was flexion-extension, with the coordinate system shown in Figure 1. Further details pertaining to setup can be found in SI1.

The patellofemoral joint was not modelled to minimise modelling assumptions in the comparative AL 288-1 specimen, in which there is no patella (note: a scaled patella is present in the AL 288-1 muscle reconstruction purely for aesthetic purposes and to assist in muscle insertion points; it is non-functional in the OpenSim model). By modelling the knee as a single, joint (no patella) fixed in position, it is acknowledged that the knee results will be simplified. The implications of this are investigated in the SI1.

Muscle masses, LoAs and segmental properties were ‘scaled’ from Subject03 (Charles et al., 2020) to TLEM2-CT via femoral length. Due to the poor quality of Subject03, it was not possible to identify discernible features of the epiphyses for precise scaling, and thus some error may have been introduced. However, the femurs from both specimens overlaid well and appeared to be of the same length. Therefore, no direct scaling was conducted, but rather a ‘re-orientation’ of Subject03’s body segments to permit all geometries to be aligned. It is acknowledged that slight error may be introduced by the impossibility of perfectly scaling these aspects, but this error is expected to be minimal and insignificant (see: Charles et al. 2020).

The specimen was set up in the ‘neutral posture’ (Wiseman et al., 2022), although in other studies these poses may differ (Bishop et al., 2021; Gatesy et al., 2022). In this pose, all joint angles were set to 0°,0°,0° (Kambic et al., 2014) from which all rotational movement deviates, with the joint axes permitting movement along each DOF. All movement deviates from this neutral posture (Wiseman et al., 2022), allowing the study and data to be comparable between subjects and reusable by future researchers.

Code provided by Bishop and colleagues (2020) was used to create a model in OpenSim 4.3 (Delp et al., 2007), permitting the computation of muscular moment arms over a range of joint motion (Charles et al., 2016; Delp et al., 2007; O’Neill et al., 2013; Seth et al., 2018; Seth et al., 2011). Each musculotendon unit (MTU) was reconstructed with reference to the muscle LoAs, as defined above. In total, this reconstruction produced 36 MTUs in the right pelvis and limb crossing the hip, knee, ankle and MTP joints – this total does not differentiate between muscles composed of multiple heads (i.e., the *M. extensor digitorum longus*). Whilst it is not currently possible to produce muscles composed of multiple heads in OpenSim (Delp et al., 2007; Hutchinson et al., 2015), multiple LoAs were created representing the same respective MTU, which were then sub-divided according to attachment site (van Beesel et al., 2021). Intrinsic muscles of the foot were not modelled; rather, only ‘foot’ muscles which crossed the ankle joint were included due to the sparsity of preserved foot material in the fossil specimen (see below). One muscle was not modelled. The *M. plantaris* is a thin, vestigial muscle, with variable attachments, often mistaken as a nerve by medical students (Spina, 2007), and missing in ∼20% of the population and also missing in 3/5 chimpanzees (Diogo et al., 2019). It is completely absent in gorillas and hylobatids (Diogo et al., 2017). Consequently, it was excluded.

*Wrapping surfaces* and *via points* were incorporated to ensure that muscle configurations were maintained throughout motion and did not penetrate through bone (Cox et al., 2019; Regnault & Pierce, 2018; Wang et al., 2004; Wiseman et al., 2021), thus improving anatomical realism of the model (Modenese & Kohout, 2020; O’Neill et al., 2013). Details regarding wrapping surface creation and visualisation, including sensitivity analyses, can be found in SI1.

To evaluate the accuracy of the human model (Hicks et al., 2015), each muscle’s moment arm was computed and compared to those primarily from the ‘gait2392 model’ (Delp et al., 1990; Yamaguchi & Zajac, 1989). Some muscle groups were subsequently found to be disparate in shape and peak timings and, to assist in model evaluation, these specific moment arms were subsequently compared to the moment arms from Charles et al. (2020) (n=10). It was found that the moment arms from this study shared an affinity with the moment arms from Charles and colleagues, with discrepancies of the gait2392 muscle attributed to small modelling differences, which are described in detail in SI1. Overall, all moment arms were considered representative of human movement, thus permitting comparative assessments between the human and AL 288-1.

### 2.2 AL 288-1 data

The freely available AL 288-1 reconstruction was used (Brassey et al., 2018), alongside other bony elements (humerus, scapula, proximal tibia, and distal femur) that were freely sourced from http://www.elucy.org. Prior to use, modifications were first made to the reconstruction of the pelvis, in which the sacroiliac and pubic joints were modified to represent improved articulation. This was deemed a necessary step owing to potential disarticulation of pelvic joints which might not reflect biological reality (Wiseman et al., 2022). Specifically, the distance between the ischiopubic ramus was reduced and both *Os coxae* were internally rotated, thus improving sacroiliac articulation whilst maintaining adequate spacing for articular cartilage (Brassey et al., 2018; Wiseman et al., 2022). All modifications were made respective to the centre of the pelvis (previously defined by Wiseman and colleagues (2022)) and thus any changes to the left side were mirrored to the right side (Figure 2).

**Figure 2.**
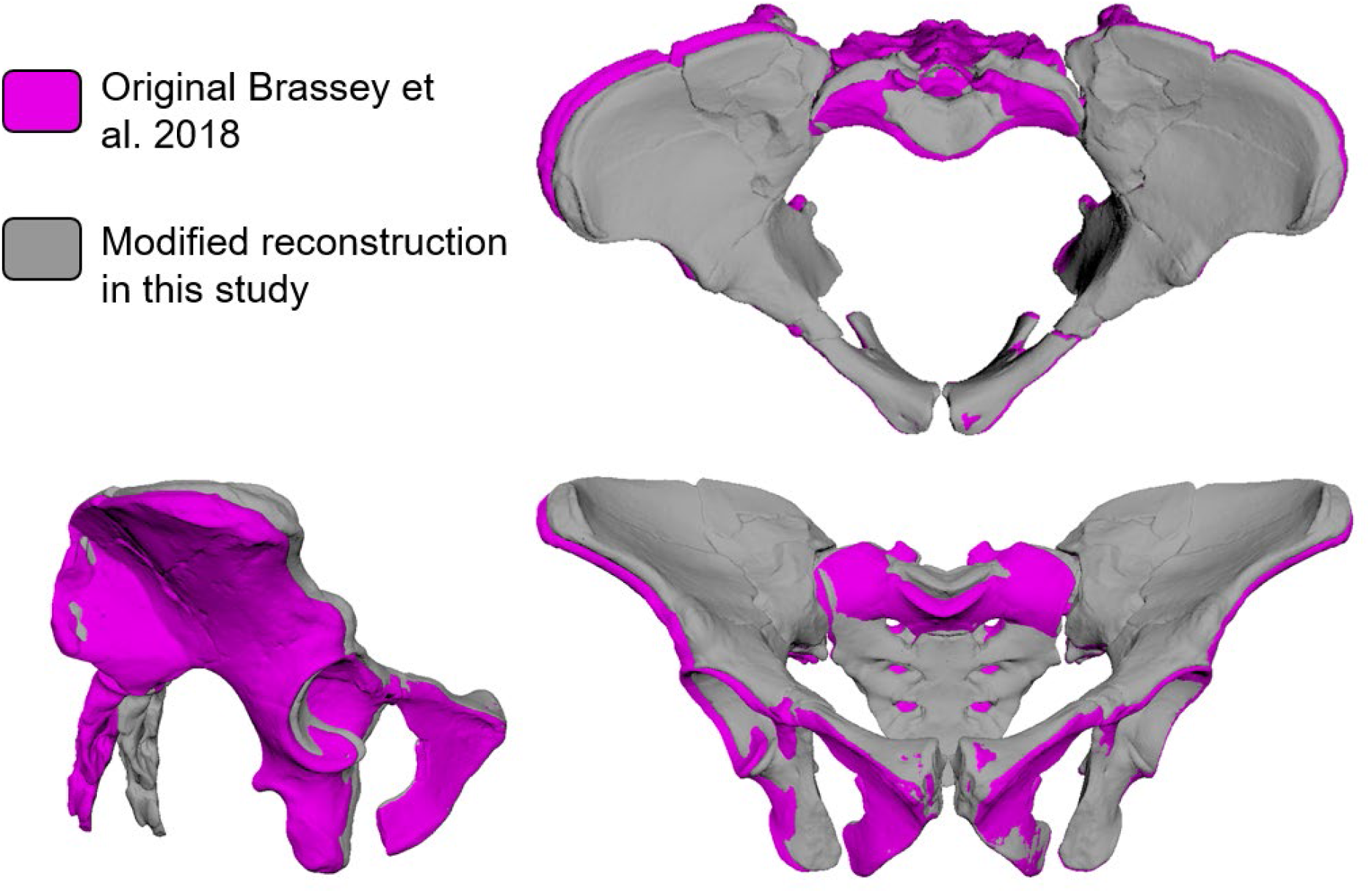
Here, the (Brassey et al., 2018) reconstructed pelvis (shown in purple) was modified (shown in grey), in which the sacroiliac and pubic joints were rearticulated. Specifically, the distance between the ischiopubic ramus was reduced and both *Os coxae* were internally rotated, thus improving sacroiliac articulation.

The tibial shaft was not previously reconstructed (Brassey et al., 2018), but was included here as a simple, non-functional cylindrical shape bridging the gap between the proximal and distal components to facilitate muscle attachments in this region. The length of the shank segment was not modified from the original reconstruction (Brassey et al., 2018), owing to previous sensitivity analyses which found that moderate changes in shank length can be ignored (Wang et al., 2004). This recreated mid-segment has no functional significance and is included here for purely aesthetic reasons.

AL 288-1 is missing a foot. A composite *Au. afarensis* foot model was created using foot bones from AL 333 scaled to the segmental lengths of AL 288-1. All available pedal remains from AL 333 were used, whilst unobtained components (the phalanges) were acquired from the human. Scaling of the phalanges to AL 333 was simplistic and subjective, and involved uniformly scaling all phalanges by the same factor associated with facet articulation. It is acknowledged that this might introduce errors into muscles which attach to the distal phalanges.

Using the same approach detailed for the human above to ensure direct comparability, a biomechanical model of AL 288-1 was created. A convex hull approach (Brassey et al., 2018; Coatham et al., 2021) was implemented in MATLAB 2021a (Mathworks, Natick) of each body segment. The produced hulls were used as a proxy for estimating body masses, centre of masses (COM) and inertial properties in OpenSim using code provided by Bishop and colleagues (2020). A correction factor was applied to each of these estimates to correct for issues in the convex hull approach which likely under-calculates segmental mass (see: Coatham et al. 2021). This approach improved all segment COM and inertia estimates (see: SI2), thus minimising potential issues with subsequent biomechanical outputs (Wang et al., 2004). Whilst COM and inertial properties do not influence moment arm calculation (see below), they were included here to standardise the manner in which musculoskeletal models of AL 288-1 (and by extent any other hominin) are constructed in the future, and also to provide the raw data for other researchers to construct a model (see below). Segmental masses were used to normalise muscle mass predictions (see below).

### 2.3 Polygonal muscle reconstructions

Obviously, it is impossible to examine muscles in a fossil via dissection or imaging techniques, and many muscle scarrings on the bones were non-identifiable. It is assumed here that the muscle attachments were most similar to that of a human rather than other analogous species owing to greater anatomical similarities (Nagano et al., 2005). However, it is acknowledged that this likely introduces some form of error and, so, the following is best classed as an ‘informed estimate’ of muscular configuration.

A 3D polygonal muscle modelling approach was implemented in Autodesk Maya 2022 (Autodesk Inc., San Rafael, USA), following the method established by Demuth and colleagues (2022). This method creates volumetric reconstructions of MTUs. Here, these reconstructions were guided by muscle scarring where visible and also by previously published cross sections of MRI scans of a human (Charles et al., 2020). For muscles in which no scarring on the bone was visible, muscle attachment was estimated based upon published dissection data from *Homo sapiens*, and loosely by data from *Pan troglodytes* (Diogo et al., 2019). To assist in this process, muscle attachment site ‘maps’ of AL 288-1 were created and are provided in SI3a. The polygonal muscle modelling approach and implementation is described in detail by (Demuth et al., 2022), but in brief:

1. Each muscle’s origin and insertion were identified and digitised. The centroid of the attachment site provided a central location for the starting point of the subsequent LoA.
2. Rows of polygon edges were extruded from each of the digitised surfaces and then united to create a singular polygonal mesh, representing a muscle’s shape.
3. The new polygonal muscle’s extent and configuration within the body was guided via cross-sectional data of humans, scaled to AL 288-1’s diaphyseal shaft diameter (see: Demuth et al., 2022). Due to pelvic morphological distinctions between a human and AL 288-1 (Lovejoy, 2005; Tague & Lovejoy, 1986), it was not possible to use cross sections in the pelvic compartment due to misalignment; instead, these were only used for the thigh and shank compartments.
4. After all muscles in a body compartment were created (i.e., all thigh musculature), remaining ‘gaps’ (i.e., spaces between the muscles) were removed by ensuring that each muscle’s belly was flush against each other, replicating biological reality. Due to morphological differences in the pelvis and limb of AL 288-1 compared with that of a human (Lovejoy, 2005, 2007), this resulted in muscle configurations that did not precisely match the cross-sections, a limitation acknowledged by Demuth and colleagues (2022). Nevertheless, the cross sections guided and informed muscular composition.
5. A LoA was automatically ‘threaded’ through each muscle’s centroid (Demuth et al., 2022), and then fed into an OpenSim 4.3 biomechanical model, following the same procedure described above for the human.

Whilst some chimpanzees have a *M. scansorius*, this is missing in humans and is instead fused with the *M. gluteus minimus* (Diogo et al., 2019; Diogo et al., 2017). Here, the human-form is assumed and was not included in the AL 288-1 model (see: Hogervorst & Vereecke (2015) for an overview of musculature differences between a human and chimpanzee).

An interactive 3D scene containing all polygonal muscles, LOAs, example cross-sectional data, ACSs and bony geometries (including the modified pelvis) is freely provided (see: SI3), in which the polygonal muscles (including their masses, configuration, origins and insertions) can be examined in detail and used in future research and/or outreach activities. Please note that the meshes of the proximal tibia and foot bones are not included.

### 2.4 Muscle mass calculations and sensitivity analyses

Muscle masses for both species were calculated assuming a standard tissue density of 1060 kg/m^3^ (Mendez & Keys, 1960) and were subsequently compared between species. Due to differences in body size, both sets of muscle masses were normalised by the respective segment mass (SI4; see: Charles et al., 2016). Notably, Subject03’s muscles are partially missing their tendons due to scan data quality. As such, their masses are likely an underestimate of their true size because only belly mass is calculable. To account for such underestimates on total muscle mass, sensitivity analyses were computed which increased each muscle’s mass by 10% and then by 15% to establish the overall influence which tendons might have on total mass – these values are greater than the 1-2 standard deviations utilised in sensitivity analyses on extant species by Demuth and colleagues (2022) and thus ensures the full spectrum of potential muscle mass under-estimation will be captured. The values here were arbitrarily selected as there is no ‘one size fits all’ ratio of tendon to belly mass for muscles. This approach was repeated for the predicted segmental masses of AL 288-1 in which the convex hull approach was applied directly to bony segments, which might have underestimated true compositional mass. The segment masses were increased by 10% and then 15% to determine how under-estimates may affect predictions on which species had relatively greater muscle mass per body segment.

### 2.5 Moment arm calculations

To test how muscle moment arms operated throughout each joint’s range of motion (flexion-extension, abduction-adduction, long-axis rotation), MTU moment arms were calculated in OpenSim 4.3 (Delp et al., 2007; Seth et al., 2011) for AL 288-1 and the human and subsequently compared. To determine if muscle moment arms peaked at the same or different limb postures between the species, moment arms per muscle group (e.g., hip extensors) were summed and then divided by the sum of the peak moment arm for each muscle, as in Hutchinson et al. (2013) and Wiseman et al. (2021). These were inspected to determine if peak moment arm values corresponded to peak limb loading postures.

The sensitivity of placing the origin and insertion of each muscle was tested by moving each attachment point up to 1 cm medio-laterally and antero-posteriorly, or proximo-distally for muscles which attached to the shafts of bones (four translations per muscle). Such method follows previous studies that tested the sensitivity of defining an attachment site in extinct species (Hutchinson et al., 2005). To assess the sensitivity of attachment sites, moment arms were calculated for each change in attachment location and compared to the original calculation. Full details are provided in the SI1.

The influence of the dimensions of a wrapping surface upon moment arm production was also assessed in which each wrapping surface was increased by ±2 cm. This value was arbitrarily selected in which increases >2 cm created wrapping surfaces located far from the bone’s surface, which produced unrealistic muscle paths. The moment arms for all muscles which crossed a particular wrapping object were plotted for comparison. Full details are provided in the SI1.

Finally, due to (1) a small sample size (n=1 fossil specimen), (2) identified moment arm variability within human models, (3) slight differences due to wrapping surface dimensions and also (3) small differences owing to attachment site location, Monte Carlo analyses were computed to provide error margins for the summed moment arms. Monte Carlo simulations were computed in MATLAB across 1000 iterations using code by Wiseman and colleagues (2021). For a given muscle, each moment arm was perturbed by a singular, randomly assigned value that produced smooth curves, rather than perturbing each timestep independent of preceding and successive timesteps. The values for each muscle were permitted to deviate up to ±20% from its original value (assuming a random uniform distribution), based upon an average of maximally varied moment arms extracted from previous studies (Brown et al., 2003; Cox et al., 2019; Wiseman et al., 2021). This method produces a ‘simulated error margin’, in which our sample data is expected to deviate within the envelope to account for the aforementioned potential issues.

## 3.0 Results

### 3.1 Pelvis modifications

The modified pelvis reconstruction differed in the following ways: there was an internal rotation of both *Os coxae*, and both the sacroiliac and pubic joints were rearticulated to minimise joint spacing (Figure 2), of which was larger in the Brassey and colleagues (2018) reconstruction than in other reconstructions (Lovejoy, 2005; Tague & Lovejoy, 1986). The new bi-acetabular diameter is 118 mm, in line with previous estimates (Tague & Lovejoy, 1986), but conflicting that of Brassey and colleagues (2018) which estimated 114.1 mm. Greater internal rotation of the *Os coxae* increased the subpubic angle from 77° (Brassey et al., 2018) to 81°, identical to the subpubic reconstruction of (Tague & Lovejoy, 1986). The modified pelvis is freely provided (SI3).

### 3.2 Polygonal muscle reconstructions

In total, 36 muscles were created per limb (Figure 3; Table 1). Sensitivity analyses were computed on each LoA’s attachment site and the influence of wrapping surface dimensions (see: SI1). The results of the sensitivity analyses demonstrated that deviations in both attachment site and wrapping surfaces are minimal, with the few differences found to be within the variability of human musculature. Therefore, without parameterisation of the muscles (not an element of the current study), biomechanical assessments should incorporate simulated error margins. Nevertheless, these reconstructions are considered the ‘best informed scenario’ with the available skeletal material.

**Table 1.** Muscles included in this study and their abbreviations. Muscles loosely ordered cranially to caudally.

| Abbreviation | Muscle name | Abbreviation | Muscle name |
| --- | --- | --- | --- |
| AB | Adductor brevis | BFL | Biceps femoris longus |
| AL | Adductor longus | BFS | Biceps femoris short head |
| AM | Adductor magnus | VI | Vastus intermedius |
| GlemInf | Glemellus inferior | VL | Vastus lateralis |
| GlemSup | Glemellus superior | VM | Vastus medialis |
| ObtExt | Obturator externus | MG | Medial gastrocnemius |
| QF | Quadratus femoris | LG | Lateral gastrocnemius |
| TFL | Tensor faciae latae | PB | Peroneus brevis |
| GMax | Gluteus maximus | PL | Peroneus longus |
| GRA | Gracilis | SOL | Soleus |
| GMed | Gluteus medius | TA | Tibialis anterior |
| GMin | Gluteus minimus | TP | Tibialis posterior |
| ILI | Iliacus | PECT | Pectineus |
| PIRI | Piriform | POP | Popliteus |
| RF | Rectus femoris | EHL | Extensor hallucis longus |
| SAR | Sartorius | FHL | Flexor hallucis longus |
| SM | Semimembranosus | EDL_DIGITS I-IV | Extensor digitorum longus |
| ST | Semitendinosus | FDL_DIGITS I-IV | Flexor digitorum longus |

**Figure 3.**
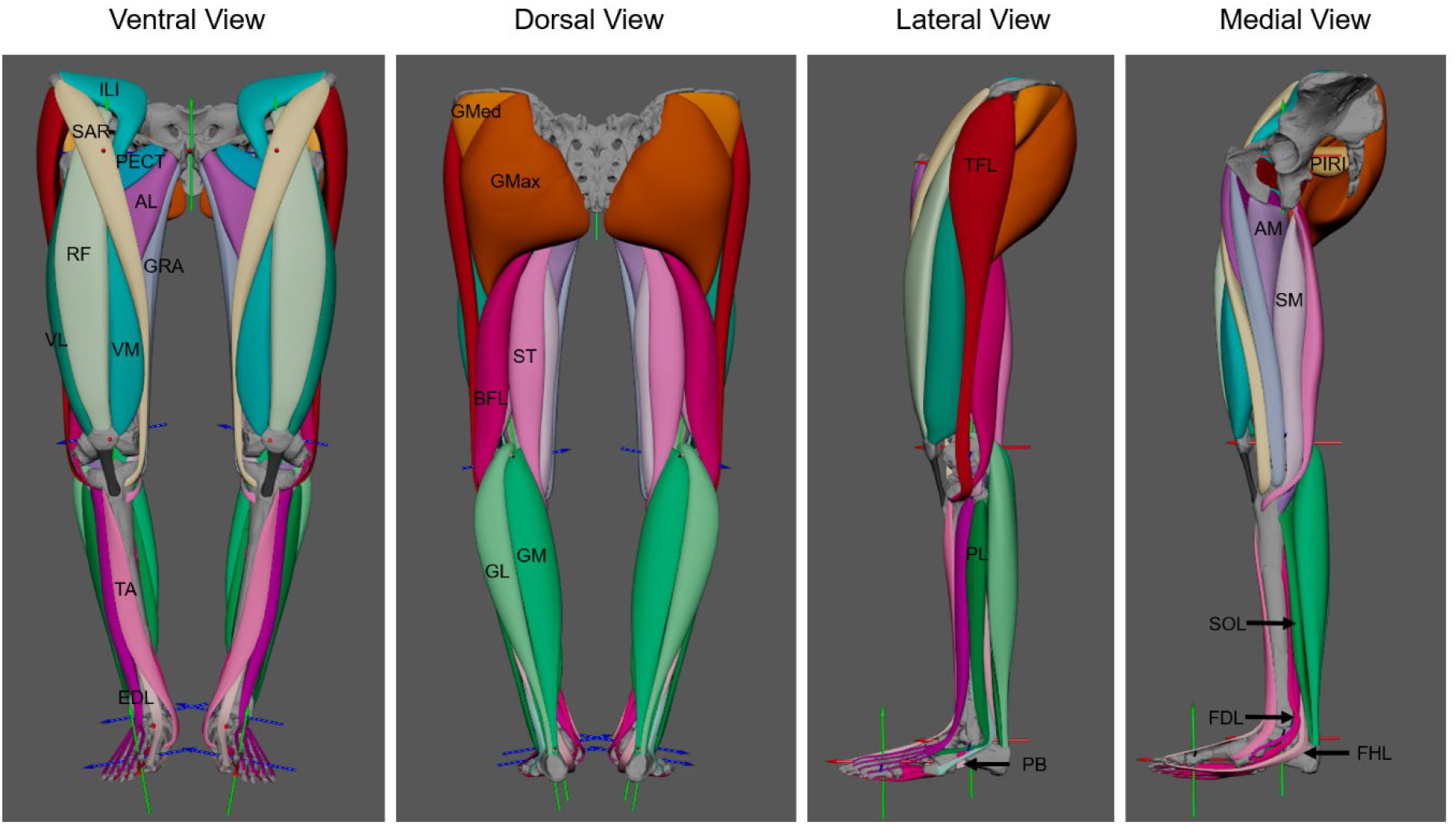
Completed views (ventral, dorsal, lateral and medial) of the polygonal muscle modelling approach, in which 36 muscles were created per lower limb – this total does not differentiate between muscles composed of multiple heads (i.e., the *M. extensor digitorum longus*). Intrinsic muscles of the foot were not modelled; rather, only ‘foot’ muscles which crossed the ankle joint were included due to the sparsity of preserved foot material. See SI2 for a diagram illustrating muscle origin and insertions in which the colours correspond to the muscle colours used here. Full muscle configurations shown here; please refer to SI3 to visualise deep musculature.

The most prominent muscular differences between AL 288-1 and a human were in the pelvic compartment. Here, increased flaring of the pelvis (Brassey et al., 2018; Lovejoy, 2005; Tague & Lovejoy, 1986) resulted in many muscles’ pelvic attachments being more transversely or frontally positioned. This had a direct impact on MTU volume and length. For example, the *M. sartorious* had a more transversely positioned origin in AL 288-1 than that of a human (the yellow polygonal muscle crossing the thigh in Figure 3; see also SI3). This muscle crosses over the front of the thigh, inserting into the superior medial tibial shaft, near the tibial tubercle. A more transversely positioned origin resulted in this muscle’s belly becoming elongated in AL 288-1 relative to a human. This muscle in humans acts as a powerful hip flexor, and differences in MTU length and volume in AL 288-1 would have likely affected function.

Other differences were found in the deep gluteal region, specifically the *Mm. glemellus inferior* and *superior* muscles (and by extent, the *Mm. quadratus femoris, piriformis* and *obturator externus*; see SI3). This external hip rotator muscle group originates from various aspects along the ischium (with the exception of *M. piriformis* which instead originates from the sacrum) and inserts into the medial aspect of the greater trochanter and trochanteric crest. AL 288-1 has a wider pelvis (Tague & Lovejoy, 1986) than a human, resulting in the distance between these muscles’ origins and insertions being relatively greater in AL 288-1 than a human. However, results of this specific muscle group should be cautiously interpreted as each muscle’s attachment site was solely inferred based upon dissection data. Poor quality MRI scan data of this compartment negated imaging guidance of the muscle path during the 3D modelling process.

The AL 288-1 pelvis was relatively shorter (Berge, 1994; Lovejoy, 2005), resulting in many muscles’ LoAs being more transversely orientated, rather than exhibiting a strong cranial orientation as in a human. For the *M. piriformis*, the LoA was almost entirely transverse in AL 288-1, rather than having an oblique angle as in a human (Charles et al., 2020). The shortness of the pelvis further affected the LoA of the *Mm. gluteus minimus* and *medius* which reduced the relative lengths of these muscles, but the wideness of the pelvis resulted in a transverse expansion of the attachment site.

Due to the configuration of the entire gluteal region (superficial and deep), it was impossible to model the *M. gluteus maximus* as having a component of its fibres attaching to the ischium, as previously suggested by Berge (1994), although this is in line with other studies (Lovejoy, 1975, 2005). This muscle must have had a human-like origin in the pelvis because the path of the *M. piriformis* prevented the *M. gluteus maximus* from occupying a space flush to the ischium, whilst other muscles’ volumes further prevented this muscle from extending deeper into the compartment. However, gluteal muscle scarring was poor and the polygonal approach was an estimate, so this should be duly noted as a potential limitation. Nevertheless, one cannot ignore other muscles’ bellies restricting gluteal expansion, and thus the reconstruction presented here is likely on-track for realism. A human-like pattern indicates potential human-like function, discussed below.

The *M. tensor fasciae latae* had a more ventro-lateral origin than in humans, possibly indicating greater capability for flexing the hip and knee. Further comparisons of this muscle would not be appropriate; this muscle varies significantly in mass and belly length/insertion in modern populations, and even in overall function (Flack et al., 2012), and thus additional comparisons are inadvisable.

AL 288-1’s femur was short relative to the pelvis in comparison to a human (Berge, 1994; Lovejoy, 1975, 2005, 2007; McHenry, 1975; Ward, 2002). Evidently, length of the quadriceps femoris muscle group was relatively shorter (knee extensors; *Mm. vastus medialis, vastus lateralis, vastus intermedius*), which originated from the *linea aspera* of the femur and inserted into the cranial surface of the patella/the quadriceps tendon into the tibial tubercle. The relatively longer human femur resulted in longer muscle lengths.

Polygonal reconstructions of the adductor group (*Mm. adductores magnus, longus, brevis*) demonstrated that these were more ventrally positioned than in humans. The attachment site of the *M. adductor magnus* was human-like owing to distinct faceting and a transverse ridge (Lovejoy, 2005) on which this muscle originated, which is a feature that is missing in non-human apes (Stern & Susman, 1983).

The AL 288-1 condyles were damaged and poorly preserved. Therefore, modelling errors may exist in muscles which wrapped around this feature (e.g., the *M. biceps femoris*). Specifically, the *Mm. popliteus, gastrocnemius lateralis* and *gastrocnemius medialis* all originated from this region and might have had the greatest source of error in the identification of attachment sites (Figure 3; see also SI3). AL 288-1 had a shorter shank than a human (Wang et al., 2004), and as such the main differences in musculature in the lower leg were due to length rather than spatial configuration.

The AL 333 foot (De Silva et al., 2018) was used as a composite to ‘complete’ the lower limb. Differences in musculature to a human (inferred from moment arms; see below) likely related to curvature of the metatarsals (Stern & Susman, 1983), morphological differences in the convexity of the cuneiform hallucal facet which likely influenced metatarsal orientation, and a less pronounced longitudinal arch (De Silva et al., 2018; Harcourt-Smith & Aiello, 2004).

### 3.3 Muscle mass comparisons

Muscle masses were compared between AL 288-1 and the human (Figure 4; SI4). Sensitivity analyses increased muscle masses in the human by 10% and by 15% to account for incomplete total muscle mass (see: Methods), but these analyses did not influence the overall trend in the results. Relative muscle masses tended to be typically smaller in AL 288-1, likely related to differences in relative segmental lengths (SI4).

**Figure 4.**
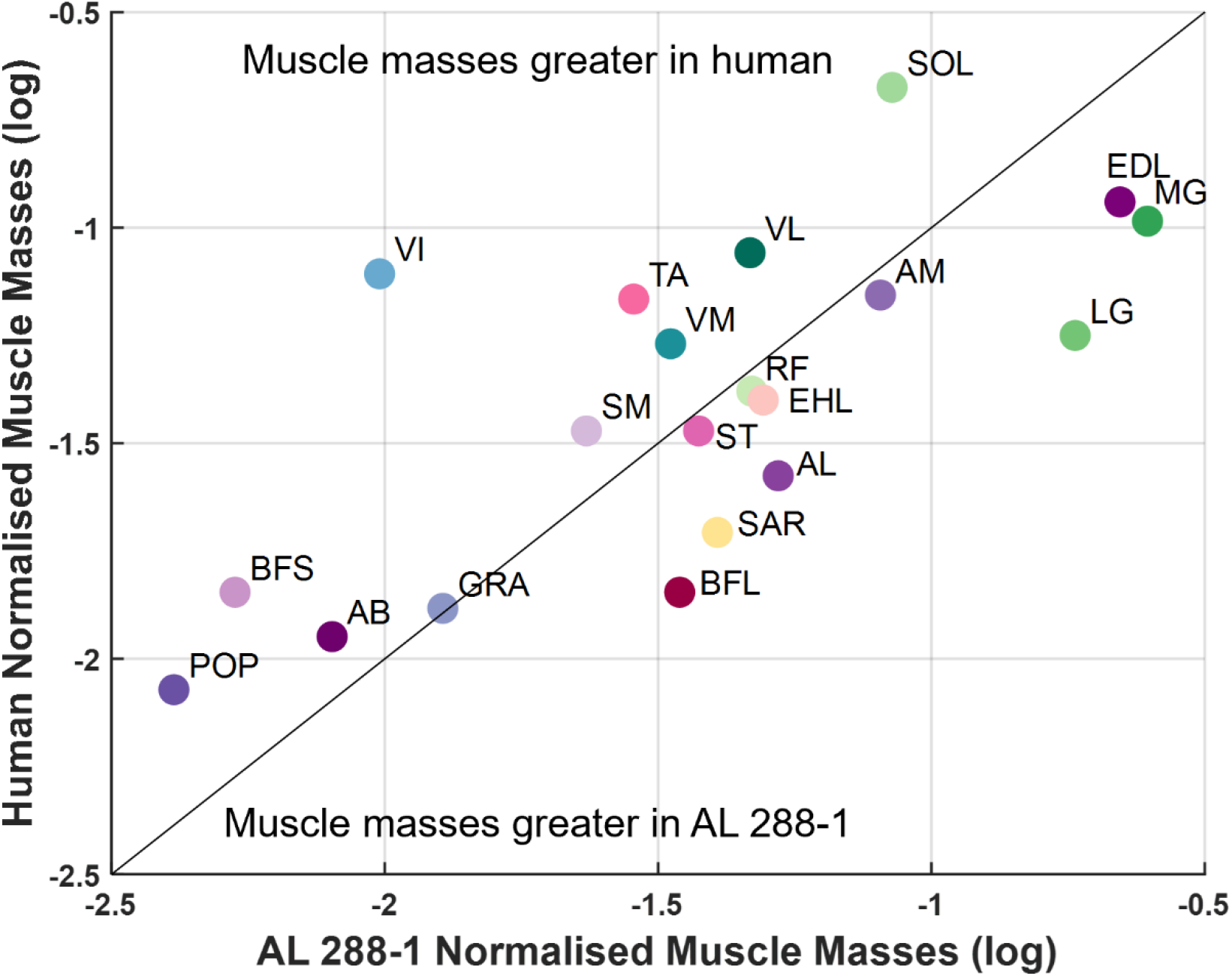
Logged normalised muscle mass calculations of AL 288-1 plotted against logged normalised muscle masses of the human. Each muscle mass was normalised by the respective segment mass in which the muscle’s belly occupies. Coloured data points correspond to the colours of muscles in Figure 3, the 3D polygonal muscles available for download in SI3, and also the muscle map diagrams in SI3a. All other muscle mass calculations for AL 288-1 can be found in SI4.

10 muscles in AL 288-1 were between 10% (*M. semitendinosus*) to 106% (*M. gastrocnemius lateralis*) greater than in the human (Figure 4), the latter of which had the greatest discrepancy by a factor of 3.3. Muscles which were smaller in AL 288-1 were mostly in the thigh compartment, in which the differences between species ranged between 2% (*M. gracilis*) and 156% (*M. vastus intermedius*). Increases in segmental and/or muscle mass (i.e., the sensitivity analyses) found that 75% of muscles were smaller in AL 288-1. If muscle masses were compared between the non-corrected segmental masses (Coatham et al., 2021), then muscle masses would have been greater in AL 288-1 (16 muscles, ranging from 22% to 167% of relative muscle muscle), in which some muscles would have been greater by a factor of 11.7 (*M. biceps femoris longus*), indicating that correction factors are indeed required for realism and uncorrected convex hull models are not sufficient and considered implausible.

### 3.4 How accurate is the model?

There are known problems and limitations with the AL 288-1 reconstruction, acknowledged by Brassey and colleagues (2018). Many components of the pelvis and limb in AL 288-1 are estimated reconstructions and the foot belongs to another specimen. Not all muscles retained evidence of muscle scarring, and thus some muscle origin and insertion sites had to be estimated (see: Methods).

Muscle masses were comparatively similar between species, implying that the predicted muscle reconstructions were anatomically realistic. If one specimen had relatively larger or smaller overall muscle mass throughout the limb, then this would have been cause for concern (Demuth et al., 2022) – but this was not the case. All configurations were guided by scaled MRI cross-sections which suggests accuracy in defining absolute segment boundaries (i.e., the configurations were not over-inflated). This was further bolstered by the comparisons of non-corrected segmental masses which found that AL 288-1 had greater relative muscle mass than a human, indicating the need for applying the correction factors to body segments. Furthermore, muscle mass in *Pan paniscus* (bonobos) exceeds that of humans in which relative muscle mass has generally decreased alongside increases in body fat composition in humans (Zihlman & Bolter, 2015). Perhaps it is unsurprising to find that 50% of muscle masses (or up to 75% if considering the sensitivity analyses) were greater in AL 288-1, a species which possibly exploited a range of habitats and locomotory repertoires and, thus, retained greater relative muscle mass than a human in some muscle groups (i.e., the hip adductors), possibly corresponding to muscle power capabilities.

Cumulatively, these factors lead to the acknowledgment that there are uncertainties and/or potential inaccuracies in this muscle modelling approach that cannot be circumvented without a full, well-preserved skeleton. Nevertheless, the reconstructed skeleton and the polygonal muscle reconstruction method provides good estimates of body composition, which can subsequently be used to infer locomotory capabilities.

Finally, it must be acknowledged that the human form was selected here as the basis for reconstructing AL 288-1’s muscles. If another primate was selected as the basis for reconstruction, the results would possibly have differed. This highlights a level of uncertainty and potential subjectivity in the reconstruction process. However, the numerous anatomical similarities between this species with a human and the many anatomical dissimilarities with a chimpanzee provide credence to our selection of the human as the baseline.

### 3.5 Moment arms

The summed moment arms for the hip flexor muscle group were greater in AL 288-1 than the human (Figure 5a). The flexor group peaked at ∼55° flexion in AL 288-1, whereas the human group peaked instead at ∼30° flexion with a lower magnitude (although, it should be reiterated that inferences based upon magnitude are somewhat redundant without associated moment generated capacity, which are not reported here). The latter corresponds to midstance postures for a range of speeds and topographies, optimising hip extension in the human for terrestrial bipedalism. The AL 288-1 peak was not observed in previously captured experimental motion data (see: Wiseman et al., 2022) and does not correspond to midstance poses. The summed moment arms for both species followed a generalised pattern, but in the human, there was a small extension ‘peak’ at ∼-35° when the limb would be positioned in an extended posture in late stance. In AL 288-1, the summed moment arms steadily increased, after which the moment arm decreased during full flexion. The similarity in overall pattern (peak timings excluded) likely elucidates comparable flexor capabilities between species.

**Figure 5.**
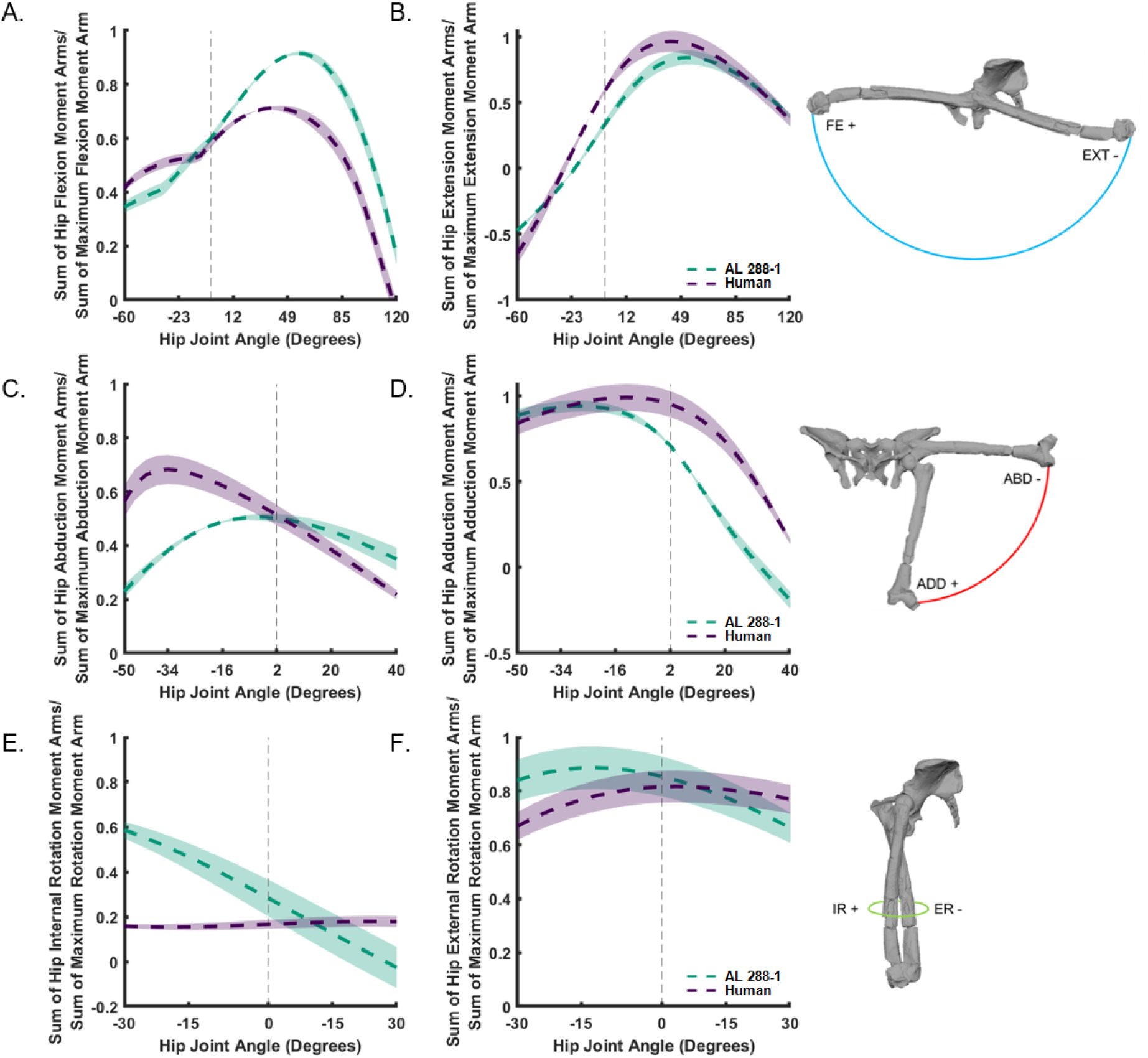
Sum of flexor (A), extensor (B), abductor (C), adductor (D), and rotator (E-F) moment arms normalised by the sum of the maximal moment arm for the associated motion, plotted against hip flexion-extension (A-B), abduction-adduction (C-D) and internal and external rotation (E-F). The horizontal grey dashed line denotes joint position 0. Example joint rotations provided.

The summed extensor moment arms were similar between species, whereby the moment arms increased as hip extension was decreased (Figure 5b). Both peaks were during hyper-extended poses of −60°. Such poses are not observed during habitual walking activities, but are still osteologically feasible.

The hip abductors were different between species (Figure 5c). In the human, the moment arms peaked at ∼43° abduction at a greater magnitude than AL 288-1 before decreasing as the hip moved into adducted poses. In AL 288-1, such changes were less pronounced with a peak at ∼-10° abduction after which the moment arm exhibited a more gradual reduction as the limb became more adducted. The hip adductors were also similarly patterned, with a gradual increase as abduction decreased and a decline during adducted poses (Figure 5d). The peak in AL 288-1 was at ∼-30° and in the human it was ∼-10°, but such variability exists within humans and is deemed inconsequential without further biomechanical data (see: SI1).

The hip internal rotator department differed in moment arm patterns and peaks (Figure 5e). The human moment arms had a near-plateau for most joint angles, with a slight increase during internally rotated postures. Contrastingly, this summed moment arm in AL 288-1 steadily decreased throughout rotation, with a peak during hyper-externally rotated postures instead. The summed moment arms of muscles assisting in external rotation were somewhat comparable between species in terms of magnitude (Figure 5f). The error margins for both groups considerably overlapped and thus it is not advisable to postulate on functional differences.

There were differences in the summed moment arms of the knee flexors (Figure 6a). The magnitude was greater in AL 288-1 during hyper-flexed postures and peaked earlier at ∼-95° flexion, before decreasing. In the human, the moment arm steadily increased during hyper-flexed postures before peaking at ∼-80° flexion which plateaued for much of knee flexion providing sustained leverage, after which there was a short reduction in moment arm length towards knee extension.

**Figure 6.**
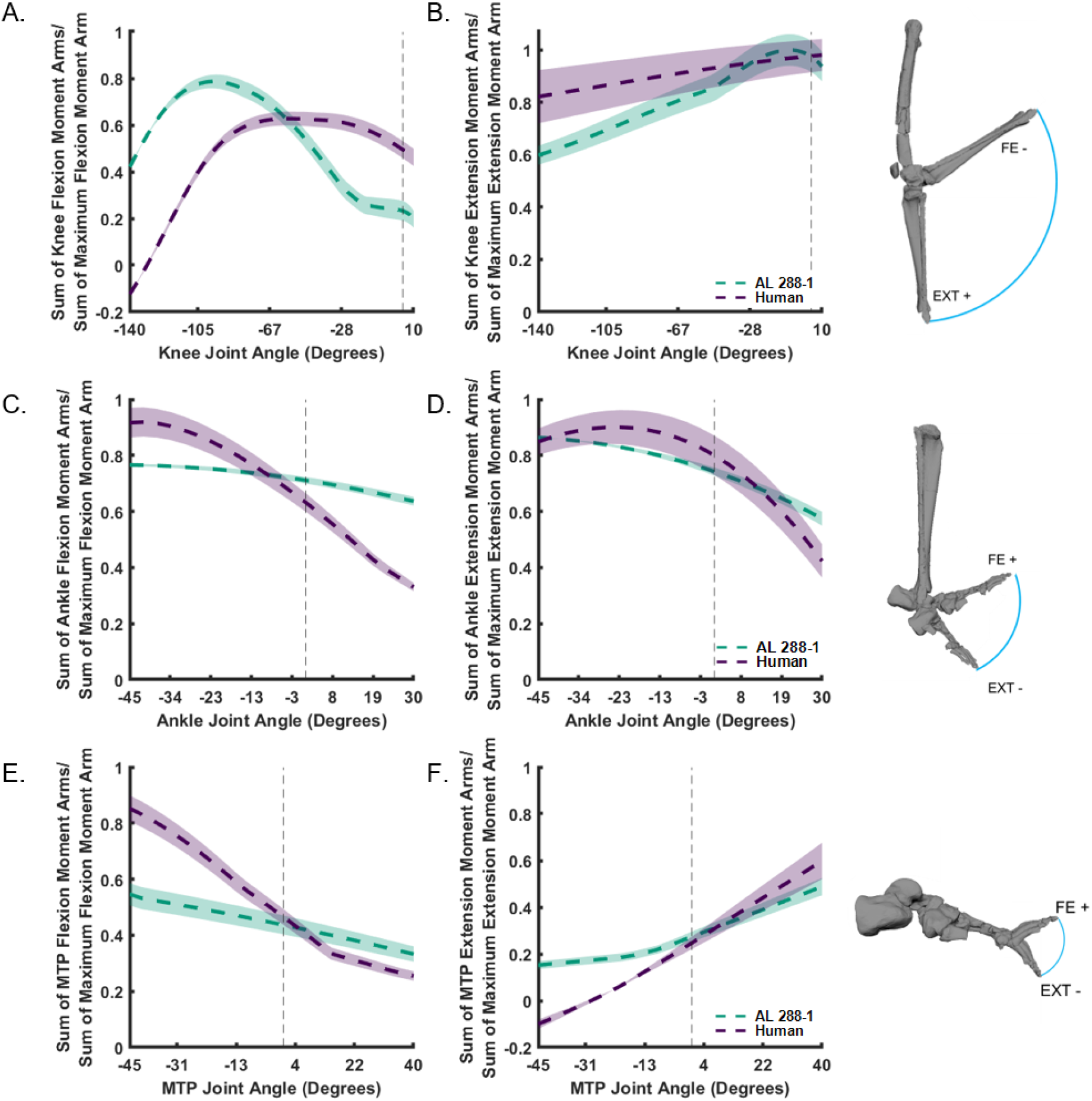
Sum of flexor (A, C, E) and extensor (B, D, F) moment arms normalised by the sum of the maximal flexor and extensor moment arms, plotted against knee flexion-extension (A-B), ankle flexion-extension (C-D) and MTP flexion-extension (E-F). The horizontal grey dashed line denotes joint position 0. Example joint rotations provided.

The summed knee extensor moment arms were similar between species. The error margins for both groups considerably overlapped towards extended postures, and thus the peak moment arms were considered the same (Figure 6b).

Both the summed moment arms for the ankle flexors and extensors had similar patterns. In AL 288-1, theses moment arm were almost at a plateau throughout the joint’s range, with a reduction in moment arm length towards dorsiflexed postures for both the flexors and extensors (Figure 6c,d). The humans had a similar pattern. Comparable patterns were also observed for the MTP flexors (Figure 6e). The summed moment arms for the MTP extensors increased during extended postures, and peaked during flexed postures (Figure 6f). The error margins overlapped and, as such, comments regarding differences in peak cannot be ascertained.

## 4.0 Discussion

This study presents a polygonal reconstruction of all musculature in the AL 288-1 pelvis and lower limb. The model calculated associated moment arms (excluding intrinsic foot musculature). These results were compared to those of a human to enhance our knowledge of the functionality of different muscle groups in the AL 288-1 limb, although it is pertinent to acknowledge that such results and inferences will be bolstered by future studies incorporating muscle parametrical data and dynamic simulations.

During walking, humans hold their hips in an adducted position during stance phase which contributes towards minimising locomotory costs during bipedal walking (Sockol et al., 2007), in contrast to chimpanzees whereby the hip is abducted during such movement (Hogervorst & Vereecke, 2015; O’Neill et al., 2015) – although a chimpanzee specimen was not included in the current study so direct comparisons cannot be made. Human hip adductors are more proximally positioned than in non-habitual bipeds (chimpanzees) to facilitate greater leverage, and differences in attachment site could, arguably, correspond to functional differences between humans and hominins. AL 288-1 exhibited a more ventrally oriented attachment site than the human but despite such anatomical differences, the summed hip adductors between them were mostly comparable. This supports previous theories that AL 288-1 was likely an efficient biped (Nagano et al., 2005; Sellers et al., 2005), and highlights that anatomical differences do not always correlate with functional differences.

However, the hip abductors tell a somewhat different story. During stance of the human gait cycle, the body’s mass is balanced over a single limb which causes rotation of the pelvis. This rotation is counteracted by powerful hip abductor muscles (Warrener et al., 2015), and so it would be expected that – if AL 288-1 was habitually an erect biped – the hip abductor group would exhibit comparable leverage to humans. The human abductors out-perform that of AL 288-1 in both magnitude and pattern. Whilst this might hint towards functional differences, similarity in other hip ranges (excluding the internal rotator compartment) suggests overall comparability in hip movement.

Obviously, anatomical differences in the pelvis and shorter segment lengths will influence muscular leverage (Brassey et al., 2017; Pandy, 1999) by impacting their length and path, but these differences do not appear to be impeding the ability to flex, extend, adduct and externally rotate the hip with respect to the human. Differences in moment arm magnitudes and peak timings could represent the repertoire of motions that AL 288-1 likely employed, perhaps ranging from bipedal walking to climbing (De Silva, 2009; Kozma et al., 2018; Stern, 2000; Venkataraman et al., 2013), but broad similarity in moment arm patterns precludes any further statement. Declarations of functional differences will only be possible with the inclusion of moment-generating capacity of which should be targeted in future studies.

The hip extensors of AL 288-1 were similar to the human which implies functional affinities. A shorter femur relative to pelvis height alongside a posterior expansion of the AL 288-1 ilium (Lovejoy, 2005; Tague & Lovejoy, 1986) optimised the gluteal compartment (specifically, the *M. gluteus maximus*) to efficiently extend the hip, thus permitting human-like bipedal leverage. This is dissimilar to the crouched bipedal posture of a chimpanzee in which this muscle is not optimised for hip extension owing to a cranial orientation of the ilium instead (Hogervorst & Vereecke, 2015; O’Neill et al., 2015; O’Neill et al., 2013; Pontzer et al., 2009; Sockol et al., 2007). This finding is in contrast to earlier studies which used simple linear action lines to estimate function (Berge, 1994), but in line with other studies that used advanced computational models (Nagano et al., 2005; Wang et al., 2004). Evidently, developments in computational methods have improved estimates of muscular leverage in extinct species, and similarity between the model presented here with previous biomechanical models (e.g., Wang et al., 2004) stresses the usefulness of polygonal modelling in advancing our knowledge of muscular configuration within the limb and of leverage.

The ischial tuberosity - where the hamstring muscle group originates (primarily knee flexors; *Mm. biceps femoris longus, semitendinosus, semimembranosus*) – has a dorsal projection in AL 288-1 relative to a human (Foster et al., 2013; Stern & Susman, 1983). Pontzer and colleagues (2009) argued that this reduces the lever arm capability of this group, favouring a habitually crouched posture rather than an extended limb in AL 288-1, although Kozma and colleagues (2018) highlighted that the leverage of this group was within the lower bounds of humans. Berge (1994) argued that the hamstrings must have had greater leverage in AL 288-1 than a human due to having a longer lever. Here, both flexor groups had a somewhat similar pattern, but different peak timings and magnitudes. The human knee is optimised for midstance postures during a period when just one limb is on the ground (Wiseman et al., 2022), but this is not the case in AL 288-1. Rather, hyper-flexed postures are optimised instead, although the moment arms are not ineffective thus indicating that the AL 288-1 knee was instead optimised for a range of movement types, beyond what we observe in humans. Overall, the results presented here are in line with Kozma and colleagues’ (2018) findings.

The knee extensors suggest that AL 288-1 was capable of efficient knee extension, which is a necessary tool for bipedal walking. Overall, the knee flexors and extensors indicate that all muscles crossing the knee were not solely suited for repetitive bipedal movement and that other locomotory behaviours were likely present and utilised. Importantly, AL 288-1 could have – and likely – walked with an erect posture of the knee (i.e., full extension), although it remains unknown if this was a regular activity.

Further down in the limb, the results do not support previous studies which argued that a shorter shank relative to humans would reduce the moment arm capability of the posterior shank compartment (*M. soleus, Mm. gastrocnemii lateralis* and *medialis*) (Wang et al., 2004). Rather, there was broad similarity in the dorsi- and plantarflexors of the ankle, which precludes any further statement as to differences in probable function. These results should be evaluated alongside the effectiveness of foot pronation and supination in the future, of which has been avoided here due to the composite AL 333 foot used and the potential for scaling issues which might influence results.

There were some differences in the flexor and extensor compartments of the foot, which could be functionally related but potential modelling errors should not be dismissed. The degree of hallucal abduction may have been insufficiently modelled here which might have influenced the results of muscles which crossed the MTP joint. On the other hand, such a result might be expected due to some possible retention of opposability (De Silva et al., 2018; Harcourt-Smith & Aiello, 2004; Holowka & Lieberman, 2018) which would have certainly affected leverage. Strong similarity in the actions of the extensor group during extension indicates similar muscular capability. This implies that AL 288-1 could have utilised the distal foot in a similar manner as a human, but further inferences are not currently possible. Future studies should incorporate forward dynamic simulations to test function with respect to applied forces (Nagano et al., 2005; Seth et al., 2011).

This study utilised a single LoA to represent each muscle as a simple demonstration of how the polygonal muscle reconstructions can be used to create musculoskeletal models. Greater fidelity in moment arm calculation will likely be achieved by creating numerous LoAs representing a singular muscle, stemming from the attachment sites – this is particularly pertinent for fan-shaped muscles, such as the gluteal muscles. Such approaches have been adopted in human biomechanical studies (Modenese & Kohout, 2020; Modenese & Renault, 2021) and might improve upon estimates of compartmental moment arms in future studies. The space-filling approach utilised here is, arguably, the best method for producing these muscle sub-divisions needed for this approach, especially for extinct taxa. Here, the polygonal muscle reconstructions provide a baseline approach for assessing muscle functionality in which all data is provided for subsequent biomechanical assessments.

### 4.1 Did AL 288-1 walk bipedally with an erect limb?

Whilst there were many key differences in muscle capability in the pelvis and limb of AL 288-1, the moment arms were overwhelmingly comparable along many joint axes, indicating that AL 288-1 was capable of an erect stance posture, as previously suggested (Foster et al., 2013; Nagano et al., 2005; Pontzer et al., 2009; Sellers et al., 2005; Vidal-Cordasco et al., 2017; Wang et al., 2004; Wiseman et al., 2022), although the efficiency and stability of such a posture should be investigated further. Differences in moment arm pattern and peak timings were expected owing to a suite of anatomical variations (Jungers, 1982), but these results support the capability of an adducted, erect limb, casting doubt upon earlier assumptions that AL 288-1 must have walked with a crouched limb posture (Stern, 2000; Stern & Susman, 1983). However, these results must be evaluated (see: Hicks et al., 2015) by future estimations of moment-generating capacity because the results presented here only tell a part of the story and cannot wholly rule out the possibility of a crouched-like posture.

Finally, the body mass of AL 288-1 is considered to be on the lower end of the range of adult body sizes for australopiths (Jungers et al., 2016). There might be implications for moment arm assessments for a smaller specimen and then subsequently extrapolating such results to the species-level, but this is impossible to ascertain without a larger australopith sample size. Nevertheless, the results here do demonstrate that AL 288-1 had similar muscular capabilities as a human, so it would seem very unlikely that larger body-sized individuals would not fall into the same category.

## 5.0 Conclusion

Here, a reconstruction of muscular configuration in the AL 288-1 pelvis and lower limb is presented, comprising of 36 muscles and their spacing within each body segment. On the basis of the results, *Au. Afarensis* was indeed capable of an erect posture, but also capable of utilising the limb in a repertoire of motions beyond habitual terrestrial bipedalism. Moving forward, the polygonal muscle modelling approach has demonstrated promise for reconstructing the soft tissues of hominins and providing information on muscle configuration and shape filling. Future studies investigating muscular function of hominins should consider (1) the composition of the body segment at question, (2) the importance of digitising the entire attachment site surface to elucidate a realistic centroid, and (3) the need to consider all muscles acting upon a body segment, not just a singular muscle (i.e., a muscle might ‘under-perform’ in one species relative to another, but other muscles might instead be contributing towards an action and, therefore, summed moment arms are preferential unless forward/inverse dynamics have also been computed in which only then singular muscle actions be investigated). An interactive 3D Autodesk Maya scene of all musculature is provided alongside this paper to aid future research and perhaps even assist in human evolutionary anatomy teaching.

## Acknowledgments

This research was supported by a Leverhulme Trust Early Career Fellowship (grant number: ECF-2021-054) and by the Isaac Newton Trust, University of Cambridge, both awarded to ALAW. Thank you to the research team associated with the reconstruction of AL 288-1 for making their model open-access (see: Brassey et al. 2018). Some bony elements were obtained from eLucy.org. Oliver Demuth is thanked for assistance in Maya Autodesk software and for providing feedback on an earlier version of this manuscript. Jeremy De Silva is thanked for providing scans of the AL 333 foot.

## Data availability statement

All data will be shared upon acceptance of the manuscript. An interactive Autodesk Maya scene will be provided open access. This scene contains all polygonal muscles, the modified pelvis, MRI cross-sections and LoAs, and the user will be able to fully explore all aspects of the reconstruction and use the models in any format for their own research (within CCBY 4.0 restrictions). Autodesk provides free academic/research licenses to all researchers/students affiliated with a university/museum/research institute. Please note that an .FBX file format is also provided which can be readily opened in other 3D modelling software (such as Blender), retaining colours, groupings and layers, although the recommended format is Maya. Whilst the FBX file can be visualised in Meshlab software, this is not recommended as colours and grouping interactability will not be accessible.

## SUPPLEMENTARY FIGURE

**Supplementary Figure 1:**
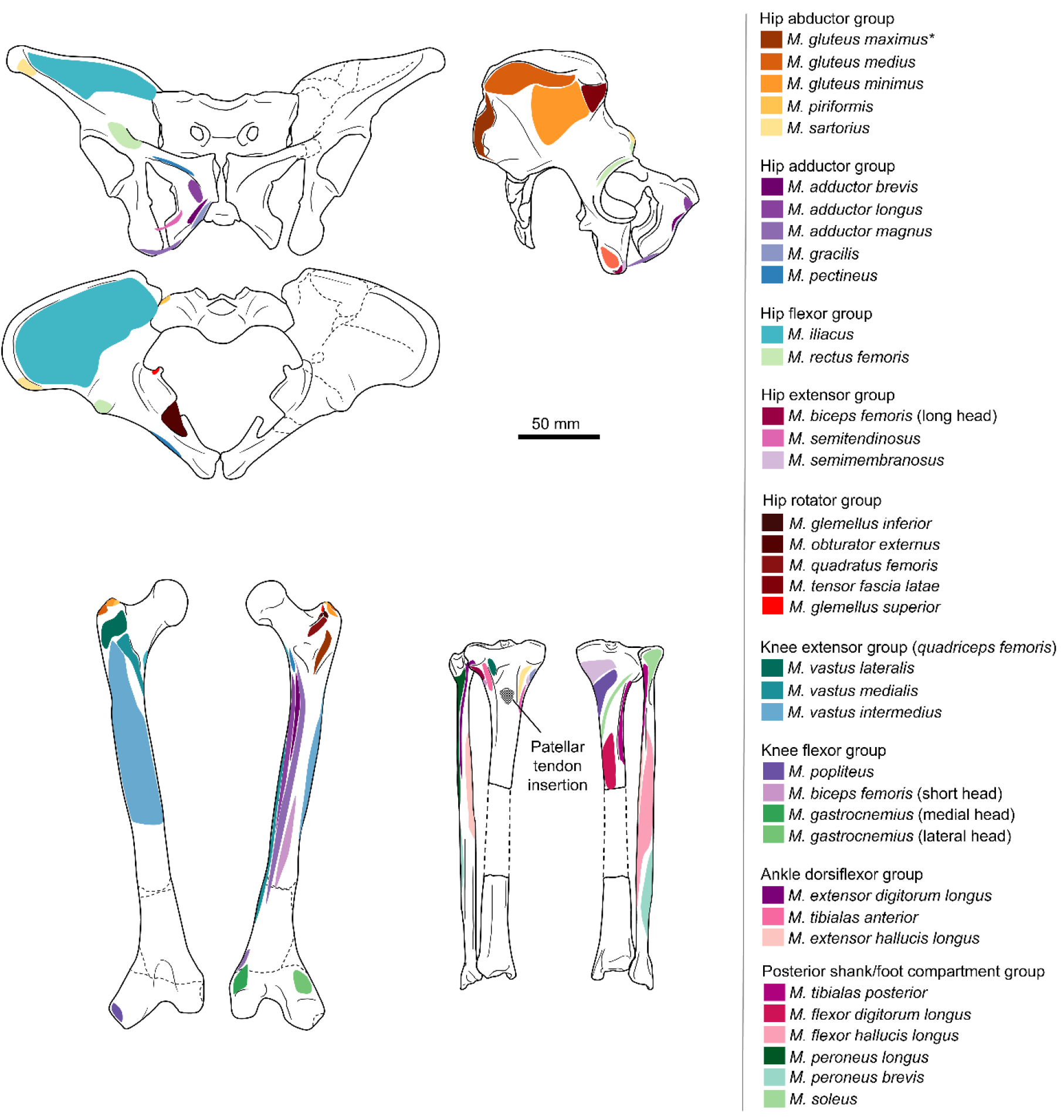
Muscle map indicating the origins and insertions of each muscle in the AL 288-1 pelvis and hindlimb.

